# Machine learning predicts early-onset acute organ failure in critically ill patients with sickle cell disease

**DOI:** 10.1101/614941

**Authors:** Akram Mohammed, Pradeep S. B. Podila, Robert L. Davis, Kenneth I. Ataga, Jane S. Hankins, Rishikesan Kamaleswaran

## Abstract

**Background:** Sickle cell disease (SCD) is a genetic disorder of the red blood cells, resulting in multiple acute and chronic complications including pain episodes, stroke, and kidney disease. Patients with SCD develop chronic organ dysfunction, which may progress to organ failure during disease exacerbations. Early detection of acute physiological deterioration leading to organ failure is not always attainable. Machine learning techniques that allow for prediction of organ failure may enable earlier identification and treatment, and potentially reduce mortality. We tested the hypothesis that machine learning physiomarkers could predict the development of organ dysfunction in an adult sample of patients with SCD admitted to intensive care units.

**Methods and Findings:** We studied 63 sequential SCD patients with 163 patient encounters, mean age 33.0±11.0 years, admitted to intensive care units, some of whom (6.7%) had pre-existing cardiovascular or kidney disease. A subset of these patient encounters (37; 23%) met sequential organ failure assessment (SOFA) criteria. The site of organ failure included: central nervous system (32), cardiovascular (11), renal (10), liver (7), respiratory (5) and coagulation (2) systems. Most (81.5%) of the patient encounters who experienced organ failure had single organ failure. The other 126 SCD patient encounters served as controls. A set of signal processing features (such as fast fourier transform, energy, continuous wavelet transform, etc.) derived from heart rate, blood pressure, and respiratory rate were identified to distinguish patients with SCD who developed acute physiological deterioration leading to organ failure, from SCD patients who did not meet the criteria. A random forest model accurately predicted organ failure up to six hours prior to onset, with a five-fold cross-validation accuracy of 94.57% (average sensitivity and specificity of 90.24% and 98.9% respectively).

**Conclusions:** This study demonstrates the viability of using machine learning to predict acute physiological deterioration heralded organ failure among hospitalized adults with SCD. The discovery of salient physiomarkers through machine learning techniques has the potential to further accelerate the development and implementation of innovative care delivery protocols and strategies for medically vulnerable patients.

## Introduction

Sickle cell disease (SCD), one of the most common genetic disorders, affects millions across the globe [1]. It was the first monogenic disorder to be characterized at the molecular level. It is characterized by the presence of abnormal hemoglobin S (HbS), which, under hypoxic conditions, causes sickling of red blood cells resulting in tissue and organ damage. Among an array of complications afflicting SCD patients, the most devastating is major organ failure, including pulmonary failure, end-stage renal disease, stroke, and heart failure [1]. A four-decade observational study reported that, by the fifth decade of life, up to half of all SCD patients had documented irreversible organ damage by the fifth decade of life [2]. Organ dysfunction may manifest or worsen during hospitalizations, when disease complications arise. Thus therapy supplemented by predictive analytics can potentially improve the potential impact of SCD on mortality, morbidity, and quality of life [3]. Early diagnosis of acute organ dysfunction may allow for early intervention, thereby preventing or reducing the severity of organ failure, particularly during hospitalization for acute complications.

In sickle cell disease, painful crises often precede the onset of acute chest syndrome, acute multiorgan failure, and other complications of the disease such as fever, altered mental status, encephalopathy, and a fall in hemoglobin and platelets [4–6]. A case review of 119 SCD patients in intensive care units identified a series of predictors, such as acute respiratory stress, and kidney injury [7]. Another study identified thrombocytopenia as an early predictor of rapidly deteriorating acute chest syndrome [6].

Early recognition of organ failure may [8,9], thereby enable clinicians to provide targeted therapies to improve outcomes. Various scoring methods have been developed for qualifying organ dysfunction, including Acute Physiology and Chronic Health Evaluation (APACHE-II) [10], Multi-Organ Dysfunction Score (MODS) [11], ‘quick’ Sequential Organ Failure Assessment (qSOFA) [12] and Sequential Organ Failure Assessment (SOFA) [13]. In this study, we used serial calculations of SOFA to identify the onset of organ failure and then used physiomarkers in machine learning models to predict organ failure for SCD patients presenting with the severe acute painful crisis. Our hypothesis was that physiomarkers [14] identified by machine learning methods can be used to predict organ failure.

## Methods and Materials

### Cohort

Continuous physiologic data was collected on 134 adult subjects with Sickle Cell Disease (comprising 318 separate patient encounters) admitted to intensive care units (ICU) at Methodist Le Bonheur Hospital (MLH), Memphis, Tennessee between June 2017 and March 2018 (Figure 1).

**Figure 1:**
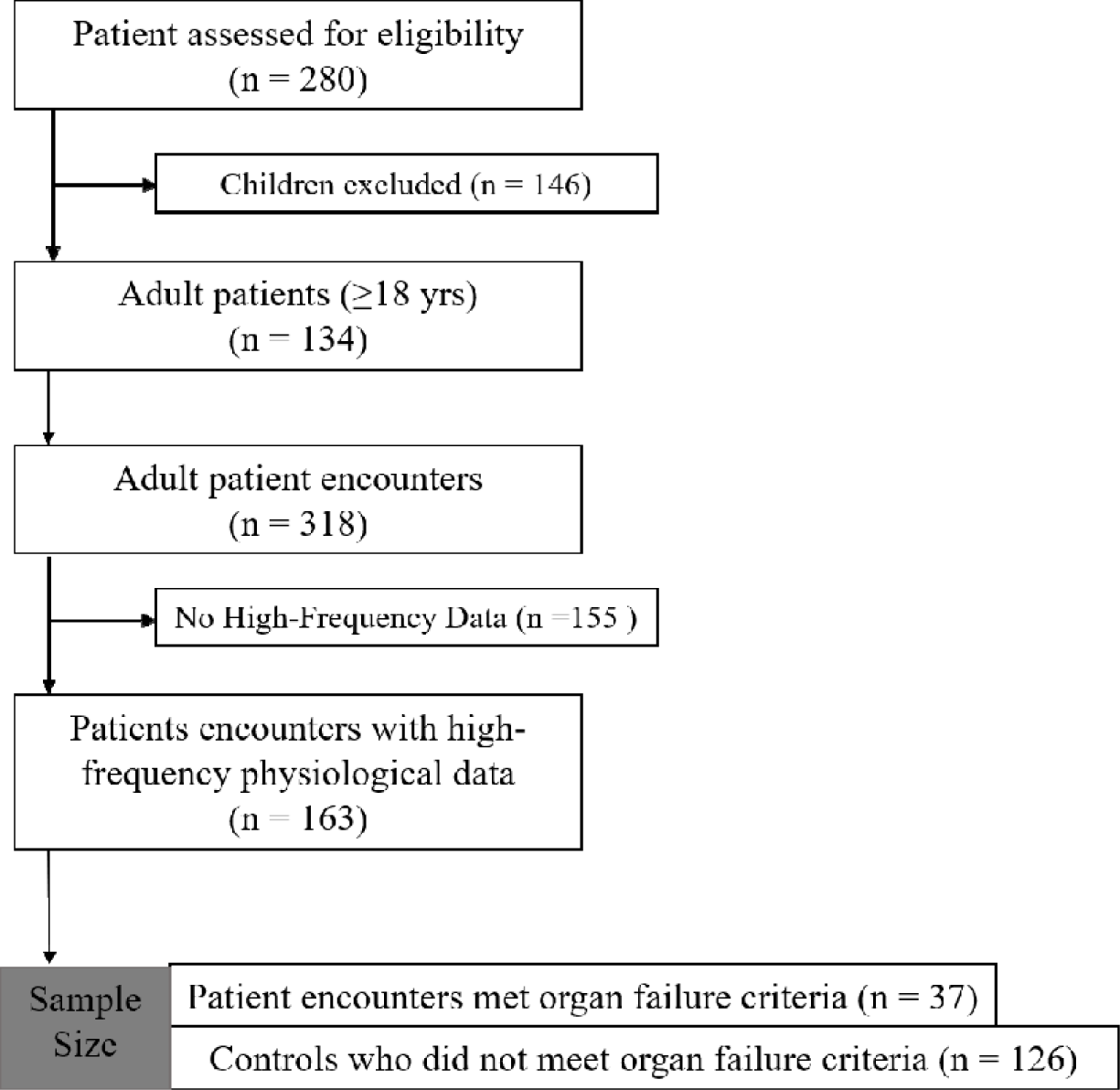
Consort diagram describing the study cohort.

All physiologic data was collected at the frequency of once per minute from the time of admission until discharge. A total of five physiologic characteristics were used in the analysis, including heart rate (HR), respiratory rate (RR), systolic blood pressure (SBP), diastolic blood pressure (DBP), and mean blood pressure (MBP). The main outcome was the development of acute physiological deterioration leading to failure of at least one organ or system (cardiovascular, liver, respiratory, coagulation, central nervous system, or renal). Consistent with the clinical definition of the SOFA score, cases were defined as patients meeting an increase in a serially calculated SOFA criteria by at least 1 score within a 24-hour rolling window, from admission till discharge. Event time (t_onset_) was recorded as the earliest timestamp of every occurence of organ failure, as defined independently by SOFA for each patient. For controls (SCD patients not developing organ failure), we extracted these same features from a 3-hour interval prior to a random time period (identified by timestamp) during their ICU stay. In order to normalize our prediction time horizon, we created relative alignments of time windows, pivoted to t_onset_. This study was approved by The University of Tennessee Health Science Center (UTHSC), and Methodist Le Bonheur Hospital Institutional Review Boards and was performed in compliance with the ethical principles for medical research involving human subjects from the Declaration of Helsinki.

### Feature Extraction and Feature Selection

We utilized Python libraries for extracting features from each of the physiological datastreams, including a combination of temporal, frequency, and statistical features (see Supplementary File 1) [15]. Features were derived from six overlapping 3-hour time intervals, with a stride of one hour, from 1-4 hours to 6-9 hours before organ failure. We performed feature selection using multiple null hypothesis testing using the Benjamini Yakutieli (BY) procedure and Mann-Whitney-U Test. Finally, we ranked the features using random forest feature importance algorithm and applied various feature thresholds to select subsets of features that most discriminatory among cases and controls.

### Machine learning algorithms

Multi-layer perceptron (MLP), support vector machine (SVM), random forest (RF), and logistic regression (LR) methods were used for building classification models. These methods were adopted because of their successful applications to medical datasets for disease classification [16–21].

A multi-layer perceptron is a deep, feed forward neural network consisting of an input layer, an output layer, and at least two or more hidden layers [22]. Non-linear activation functions are integrated into each neuron within the hidden layers for extracting meaningful relationships between various input features. MLP has been used in a variety of applications, including EEG signal classification [18], heart disease diagnosis [23], ovarian tumor classification [24], and continuous speech recognition [25].

The multi-layer perceptron architecture used in this study consisted of 5 hidden layers, with the number of neurons decreasing after each layer. The five layers consisted of 512, 256, 128, 64, and 16 neurons respectively. We applied batch normalization [26], before activation using the rectified linear unit. To avoid overfitting, we further imposed a dropout [27] ratio of 0.3. The output layer performed binary classification using the sigmoid activation, and our loss function used the Adam optimizer.

The support vector machine is a multivariate machine learning approach for classifying samples through pattern recognition analysis [28]. SVM aims to find the best hyperplane that separates all data points of one class compared to those of another class. We used a radial basis function as the kernel parameter for model building.

The random forests classifier is well suited to the classification of medical data because of the following advantages: (i) it performs embedded feature selection (ii) it incorporates interactions between predictors; (iii) it allows the algorithm to learn both simple and complex classification functions accurately; and (iv) it is applicable to both binary and multicategory classification tasks [29]. Based on the out-of-bag error [30], we identified 500 trees in the RF models as the optimal number of trees.

Logistic regression can be used as a machine learning method used to predict the value of a binary variable based on its relationship with predictor variables [31]. The p-value for statistical testing of variable significance for *inclusion-in* and *exclusion-from* the model was set to 0.05 and 0.10 respectively, and liblinear solver was used for the optimization function.

### Statistical Analysis and machine learning framework

Python scikit-learn machine learning library [32] was used for calculating descriptive statistical measures, feature selection and building machine learning classifiers. Bivariate logistic regression, bootstrap and Bayesian bootstrap (adjusted for weights) were used to assess the predictability of the features generated for predicting organ failure [33].

### Predictive Performance Measures

Accuracy was defined as the ability of models to categorize testing sample data correctly. Accuracy, specificity, and sensitivity are derived from the measures mentioned above; accuracy is the ratio of correctly predicted samples to the total number of samples. Specificity is the proportion of true negatives which are predicted as negatives whereas, sensitivity is the proportion of true positives that are predicted as positive.

### Cross-Validation

Models were developed from the distinct time intervals and tested on patients who were not included in the training of the model. For model selection and accuracy estimation, we used 5-fold cross-validation [34]. This technique divides data (stratified partitioning) into five equal and discrete folds and uses four folds for the model generation, while predictions are generated and evaluated using the remaining one-fold. This step is subsequently repeated five times, so each fold is tested against the other four folds. We further ran each of these 5-fold cross-validation models ten times by shuffling the data in each iteration and averaged the performance metrics from all iterations to reduce bias.

## Results

### Patient characteristics

Table 1 and 2 outline the descriptive level characteristics of 63 patients and clinical characteristics of 163 encounters respectively.

**Table 1.**
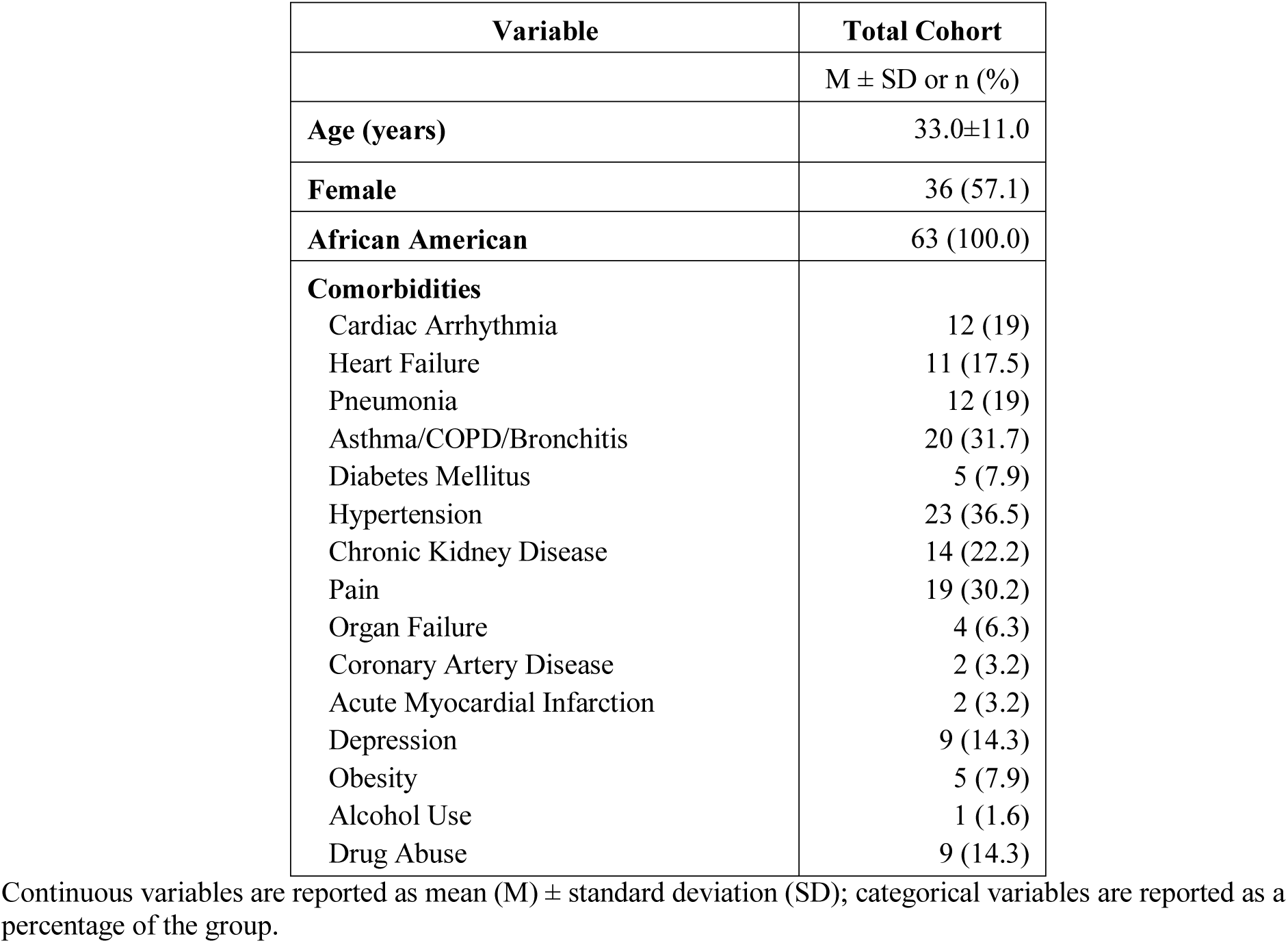
Demographic and comorbidities of patients in the overall cohort (n=63).

| Variable | Total Cohort |
| --- | --- |
|  | M ± SD or n (%) |
| Age (years) | 33.0±11.0 |
| Female | 36 (57.1) |
| African American | 63 (100.0) |
| **Comorbidities** |  |
| Cardiac Arrhythmia | 12 (19) |
| Heart Failure | 11 (17.5) |
| Pneumonia | 12 (19) |
| Asthma/COPD/Bronchitis | 20 (31.7) |
| Diabetes Mellitus | 5 (7.9) |
| Hypertension | 23 (36.5) |
| Chronic Kidney Disease | 14 (22.2) |
| Pain | 19 (30.2) |
| Organ Failure | 4 (6.3) |
| Coronary Artery Disease | 2 (3.2) |
| Acute Myocardial Infarction | 2 (3.2) |
| Depression | 9 (14.3) |
| Obesity | 5 (7.9) |
| Alcohol Use | 1 (1.6) |
| Drug Abuse | 9 (14.3) |

Continuous variables are reported as mean (M) ± standard deviation (SD); categorical variables are reported as a percentage of the group.

**Table 2.** Encounter level clinical characteristics of patients in the overall cohort (n=163).

| Variable | Total Cohort | Organ failure (Yes) | Organ failure (No) | *p-value* |
| --- | --- | --- | --- | --- |
|  | M ± SD or n (%) | M ± SD or n (%) | M ± SD or n (%) |  |
| **Number of encounters** | **163 (100.0%)** | **37 (22.7%)** | **126 (77.3%)** |  |
| **Encounter through ED** | 134 (82.2) | 33 (89.2) | 101 (80.2) | 0.33 |
| **Length of stay (days)** | 5.3±4.7 | 7.7±8.4 | 4.5±2.5 | 0.03* |
| **APR-DRG Severity of Illness (SOI)** |  |  |  | <0.0001* |
| Minor | 52 (37.1) | 7 (19.4) | 45 (43.2) |  |
| Moderate | 40 (28.6) | 7 (19.4) | 33 (31.7) |  |
| Major | 40 (28.6) | 15 (41.7) | 25 (24.0) |  |
| Extreme | 8 (5.7) | 7 (19.4) | 1 (1.0) |  |
| **APR-DRG Risk of Mortality (ROM)** |  |  |  | <0.0001* |
| Minor | 93 (66.4) | 14 (38.9) | 79 (76.0) |  |
| Moderate | 26 (18.6) | 6 (16.7) | 20 (19.2) |  |
| Major | 12 (8.6) | 8 (22.2) | 4 (3.9) |  |
| Extreme | 9 (6.4) | 8 (22.2) | 1 (1.0) |  |
| **Charlson Comorbidity Index (CCI)** |  |  |  | 0.006* |
| 0 | 107 (65.6) | 17 (46.0) | 90 (71.4) |  |
| 1 | 29 (17.8) | 6 (16.2) | 23 (18.3) |  |
| 2 | 9 (5.5) | 3 (8.1) | 6 (4.8) |  |
| ≥3 | 18 (11.0) | 11 (29.7) | 7 (5.6) |  |
| **Discharge Disposition** |  |  |  | <0.0001* |
| Home | 147 (90.2) | 24 (64.9) | 123 (97.6) |  |
| Hospice/Home Health Services | 7 (4.3) | 5 (13.5) | 2 (1.6) |  |
| Expired | 5 (3.1) | 5 (13.5) | 0 (0.0) |  |
| Other | 4 (2.5) | 3 (8.1) | 1 (0.8) |  |
| **Discharge Diagnosis** |  |  |  | 0.0005* |
| SCD with Crisis (All genotypes) | 111 (41.7) | 15 (22.2) | 96 (47.3) |  |
| SCD with ACS | 9 (6.7) | 3 (0.0) | 6 (8.6) |  |
| Sepsis | 10 (9.2) | 6 (25.9) | 4 (4.3) |  |
| Organ Failure (Heart/Kidney) | 10 (6.7) | 5 (14.8) | 5 (4.3) |  |
| Pneumonia | 5 (5.0) | 1 (3.7) | 4 (5.4) |  |
| Other | 18 (27.5) | 7 (29.6) | 11 (26.9) |  |

Continuous variables are reported as mean (M) ± standard deviation (SD); categorical variables are reported as a percentage of the group. APR-DRG: all patient refined-diagnosis related group. For continuous variables independent t-test was used, and for categorical variables, chi-square test of independence was used. Fisher's exact test was used for variables with cell counts of less than 5.

The mean age of the patients in the cohort was 33.0±11.0 years, all patients were African American, and there were more females (36; 57.1%) than males (27; 42.9%). Comorbidities were common, and included comorbidities of hypertension (36.5%), acute chest syndrome (33.3%), asthma (31.7%), pain (30.2%), and chronic kidney disease (22.2%) (Table 1). Patients with organ failure had longer hospital stays (3.2 additional hospital days, p=0.03) than controls, and had higher severity of illness (p <0.0001), and risk of mortality (p <0.0001). Charlson comorbidity index (CCI) score of ≥3 (p =0.006) (Table 2). There were a total of 67 episodes of organ failure occurring among the 37 cases. The central nervous system (i.e. encephalopathy) was the most frequently reported involved (32), followed by heart failure (11), acute kidney failure (10), renal failure (7), and respiratory failure (5).

### Feature selection

Feature selection was performed to reduce the number of features, and the reduced feature set was fed into each of the classifiers. The sample distribution for each of the six datasets for 3-hour observational periods is given in Table 3. The number of patients varies with the availability of data during each time window.

**Table 3:**
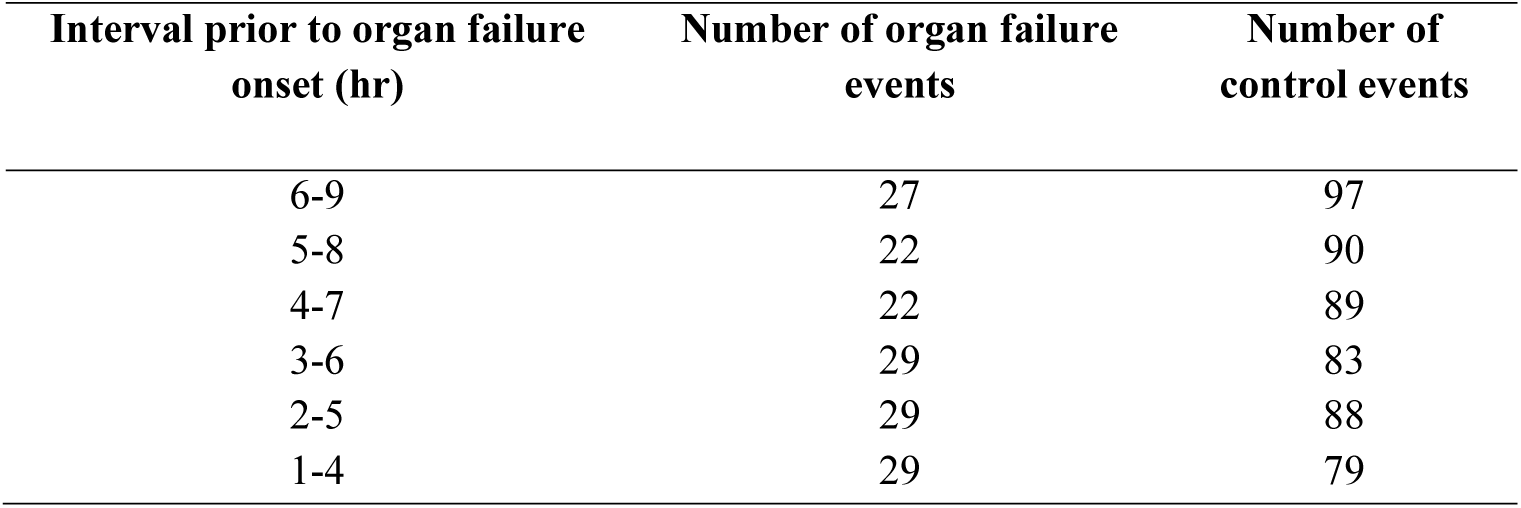
Sample distribution and number of features for each dataset using organ failure

### Model performance

Five-fold cross-validation models were developed using multi-layer perceptron, random forests, support vector machine, and logistic regression for each of the six datasets. The average sensitivity and specificity from all models for each of the six time-periods are given in Figure 2. The RF model achieved an overall 5-fold cross-validation accuracy of 94.57% with average sensitivity and specificity of 90.24% and 98.9% respectively, 6 hours before organ failure (Table 4 and Figure 2). Among the four classifiers, the RF performed better than support vector machine, logistic regression, and multi-layer perceptron in predicting SCD with organ failure.

**Table 4:** Average 5-fold Cross-Validation Accuracies of SVM, RF, LR, and MLP models. SVM: support vector machine, RF: random forest, LR: logistic regression, MLP: Multi-layer perceptron, CI: confidence interval

| Time before organ failure (hour) | Interval analyzed before organ failure (hour) | SVM mean accuracy [95% CI] | RF mean accuracy [95% CI] | LR mean accuracy [95% CI] | MLP mean accuracy [95% CI] |
| --- | --- | --- | --- | --- | --- |
| 1 | 1-4 | 89.36 [87.03, 91.69] | 96.66 [95.11, 98.21] | 88.51 [85.77, 91.25] | 89.57 [86.33, 92.81] |
| 2 | 2-5 | 93.75 [92.05, 95.44] | 95.15 [93.71, 96.59] | 95.09 [93.73, 96.45] | 92.01 [89.31, 94.71] |
| 3 | 3-6 | 94.15 [92.31, 95.99] | 94.03 [92.04, 96.02] | 95.27 [93.39, 97.15] | 89.64 [86.44, 92.84] |
| 4 | 4-7 | 90.45 [87.70, 93.20] | 96.27 [94.34, 98.20] | 88.5 [86.03, 90.97] | 93.80 [90.96, 96.64] |
| 5 | 5-8 | 94.22 [92.03, 96.41] | 95.59 [93.57, 97.61] | 96.45 [95.00, 97.90] | 92.64 [89.61, 95.67] |
| 6 | 6-9 | 93.27 [91.71, 94.83] | 94.57 [93.03, 96.11] | 94.72 [92.92, 96.52] | 89.88 [87.11, 92.65] |

**Figure 2:**
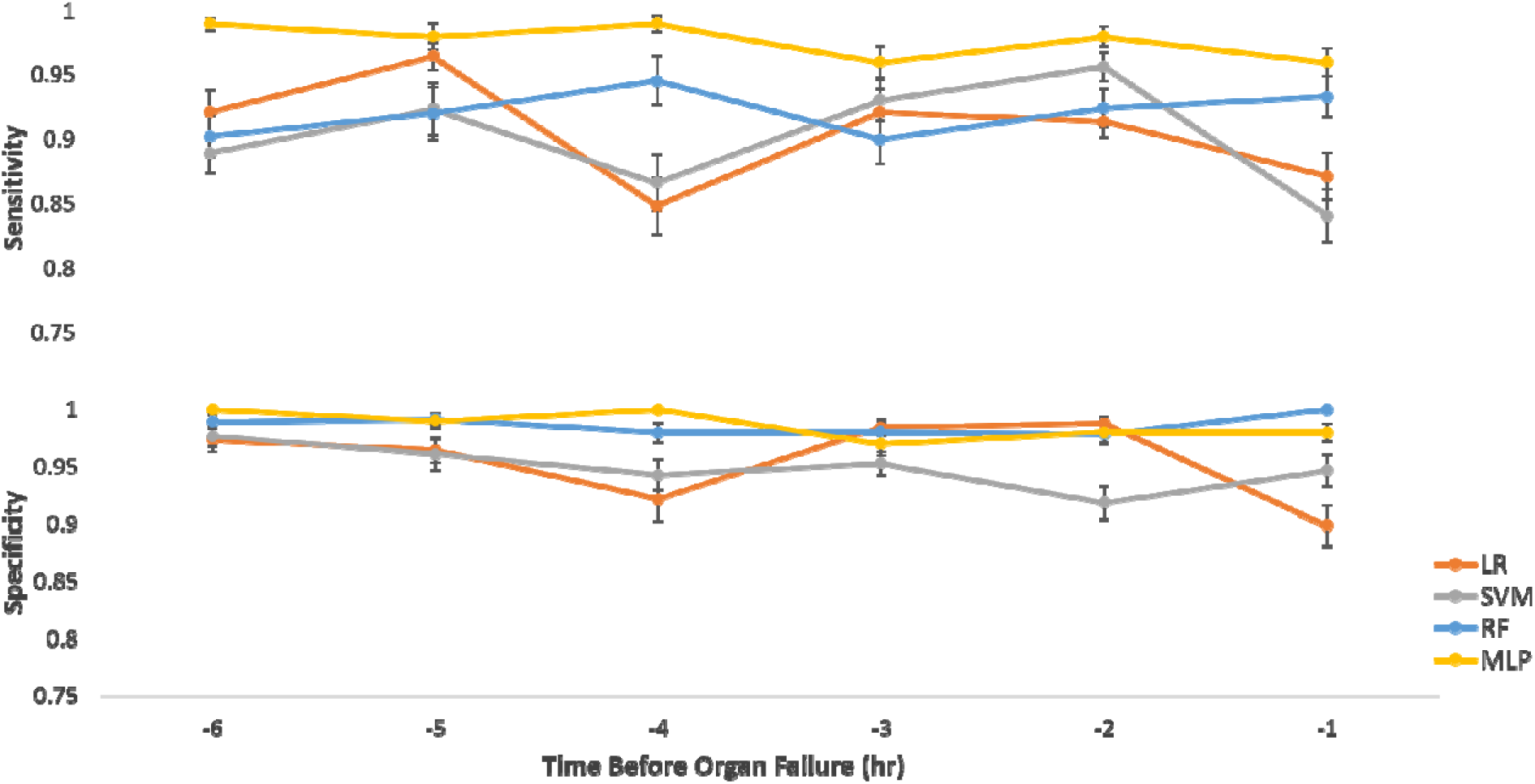
Average sensitivity, specificity for support vector machine (SVM), random forest (RF), logistic regression (LR), multi-layer perceptron (MLP) sickle cell disease models using each of the six 3-hour datasets.

We extracted feature importance from each of the selected feature sets using random forest classifier (random forest provides an option of pruning trees below a particular impurity (gini impurity) value at the end-level of the trees and rank the most important feature by how well it improves the purity of the node in a tree) and identified the continuous wavelet transform generated from mean blood pressure time-series physiologic variable as the most important feature, followed by continuous wavelet transform feature generated from respiratory rate of the time-series data. Figure 3 shows the frequency of the top 30 important features generated from five physiologic signals that are used to build the machine learning models. Each box in the heatmap represents the frequency of a feature (y-axis) generated from the physiological signal (x-axis). The dark purple color in the figure 3 represents the absence of the feature whereas dark yellow color represents the most frequently present feature. The list of features ranked by their importance is given in Supplementary File 2. The description of each feature is presented in Supplementary File 1.

**Figure 3:**
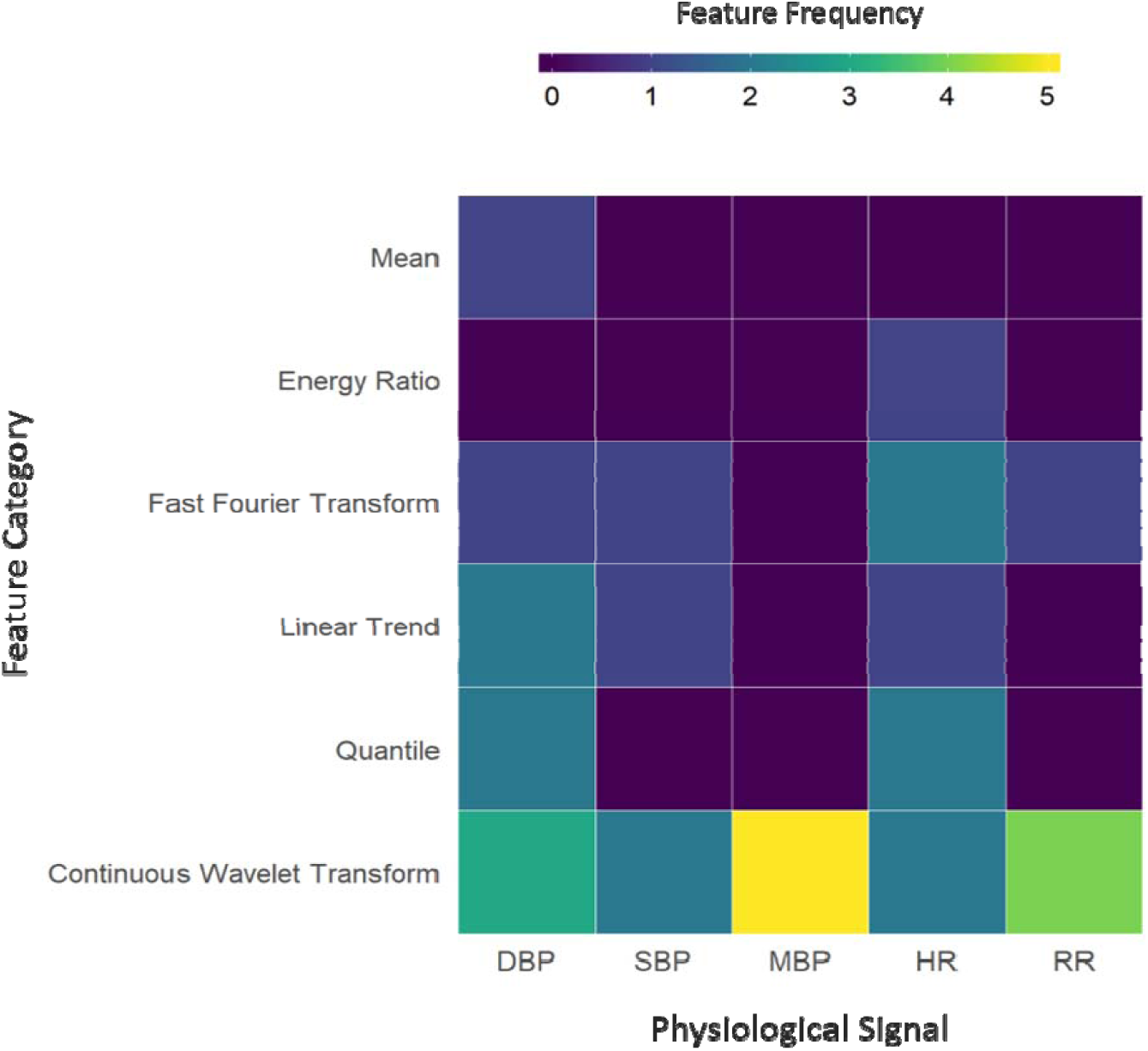
Features derived from physiologic signals up to six hours prior to organ failure. DBP = diastolic blood pressure, SBP=systolic blood pressure, MBP= mean blood pressure, HR= heart rate, RR= respiratory rate

## Discussion and Conclusion

Acute organ failure is a major challenge in persons with sickle cell disease, especially among adults experiencing an acute disease complication. The ability of predictive algorithms to identify patients at high risk for organ deterioration by using routinely collected physiological data can provide important early warnings of impending physiological deterioration. Such information can aid clinical decision making and may even eventually be useful in guiding early goal-directed therapy. In this study, we demonstrated that the machine learning-based prediction models could accurately distinguish SCD patients at risk for developing organ failure up to 6 hours prior to the onset. The classifiers and the selected physiologic features may facilitate accurate, unbiased SCD diagnosis and effective treatment, ultimately improving prognosis.

The selection of relevant features involved in SCD with organ failure and control remains a challenge [35,36]. Therefore, it is important to find a subset of physiologic features that are sufficiently informative to distinguish between SCD patients at-risk of developing organ failure and those who are not. To extract useful information from SCD patients’ continuous physiologic data and to reduce dimensionality, feature-selection algorithms were systematically investigated in this study. As we have demonstrated in the results, selecting smaller subsets of features allowed for the high performance of our classification models. Salient physiomarkers (such as fast fourier transform, energy, continuous wavelet transform, etc.) derived from the physiological signals such as blood pressure, heart rate, and respiratory rate, may precede acute organ failure in SCD patients as suggested by the results in this study. A shortcoming of machine learning is that these physiomarkers are neither observable by physicians nor are they readily interpretable; instead their benefit here is primarily towards the early prediction of impending physiologic deterioration and alerting health care providers of that fact. Further research is needed to understand how to use these alerts to guide personalized care of patients with SCD.

Other limitations are also important to mention. First, we developed the machine learning model on a small subset of patients, specific to the Mid-South USA, potentially reducing generalizability. Moreover, the data was highly imbalanced, with more non-organ failure cases compared to organ failure cases, making data-driven approaches difficult to implement. Missing data elements identified in the data was a major hindrance for model validation, and thus may have contributed to poor validation in some of the cross-validation folds. Therefore, additional data are required to develop a more robust and generalizable model. The development of such a model can greatly improve the prediction of SCD patient outcomes, including potentially a reduction in mortality and morbidity associated with such rapid deteriorations.

In conclusion, we showed, as a proof of principle, that machine learning can accurately predict the development of organ failure in ICU patients with SCD up to 6 hours before onset. This finding is significant because it may optimize the early recognition of serious disease complications and allow for the implementation of early interventions.

## Acknowledgments

We would like to acknowledge Brian Williams, Michael Younker, and Don MacMillan for their assistance in the data collection.

## References

1. Powars DR. Sickle cell anemia and major organ failure. Hemoglobin. 1990;14:573–598. doi:10.3109/03630269009046967

2. Thein MS, Igbineweka NE, Thein SL. Sickle cell disease in the older adult. Pathology. 2017;49: 1–9. doi:10.1016/j.pathol.2016.10.002

3. Powars DR, Chan LS, Hiti A, Ramicone E, Johnson C. Outcome of sickle cell anemia: A 4-decade observational study of 1056 patients. Medicine (Baltimore). 2005;84: 363–376. doi:10.1097/01.md.0000189089.45003.52

4. Hassell KL, Eckman JR, Lane PA. Acute multiorgan failure syndrome: A potentially catastrophic complication of severe sickle cell pain episodes. Am J Med. 1994;96: 155–162. doi:10.1016/0002-9343(94)90136-8

5. Novelli EM, Gladwin MT. Crises in sickle cell disease. Chest. 2016;149: 1082–1093. doi:10.1016/j.chest.2015.12.016

6. Chaturvedi S, Ghafuri DL, Glassberg J, Kassim AA, Rodeghier M, DeBaun MR. Rapidly progressive acute chest syndrome in individuals with sickle cell anemia: a distinct acute chest syndrome phenotype. Am J Hematol. 2016;91: 1185–1190. doi:10.1002/ajh.24539

7. Cecchini J, Lionnet F, Djibré M, Parrot A, Stojanovic KS, Girot R, et al. Outcomes of adult patients with sickle cell disease admitted to the ICU: A case series. Crit Care Med. 2014;42: 1629–1639. doi:10.1097/CCM.0000000000000316

8. Johnson AEW, Ghassemi MM, Nemati S, Niehaus KE, Clifton D, Clifford GD. Machine Learning and Decision Support in Critical Care. Proc IEEE. 2016;104: 444–466. doi:10.1109/JPROC.2015.2501978

9. Kamaleswaran R, Akbilgic O, Hallman MA, West AN, Davis RL, Shah SH. Applying Artificial Intelligence to Identify Physiomarkers Predicting Severe Sepsis in the PICU. Pediatr Crit Care Med. United States; 2018;19: e495–e503. doi:10.1097/PCC.0000000000001666

10. Knaus WA, Draper EA, Wagner DP, Zimmerman JE. APACHE II: A severity of disease classification system. Crit Care Med. 1985;13: 818–829. doi:10.1097/00003246-198510000-00009

11. Marshall JC, Cook DJ, Christou N V., Bernard GR, Sprung CL, Sibbald WJ. Multiple organ dysfunction score: A reliable descriptor of a complex clinical outcome. Critical Care Medicine. 1995. pp. 1638–1652. doi:10.1097/00003246-199510000-00007

12. Singer M, Deutschman CS, Seymour C, Shankar-Hari M, Annane D, Bauer M, et al. The third international consensus definitions for sepsis and septic shock (sepsis-3). JAMA - J Am Med Assoc. American Medical Association; 2016;315: 801–810. doi:10.1001/jama.2016.0287

13. Vincent J-L, Moreno R, Takala J, Willatts S, Mendona A De, Bruining H, et al. The SOFA (Sepsis-related Organ Failure Assessment) score to describe organ dysfunction/failure. Intensive Care Med. 1996;22: 707–710. doi:10.1007/BF01709751

14. van Wyk F, Khojandi A, Mohammed A, Begoli E, Davis RL, Kamaleswaran R. A minimal set of physiomarkers in continuous high frequency data streams predict adult sepsis onset earlier. Int J Med Inform. 2019;122: 55–62. doi:10.1016/j.ijmedinf.2018.12.002

15. Christ M, Braun N, Neuffer J, AW Kempa-Liehr. Time Series FeatuRe Extraction on basis of Scalable Hypothesis tests (tsfresh – A Python package). Neurocomputing. Elsevier; 2018;307: 72–77. doi:10.1016/j.neucom.2018.03.067

16. Statnikov A, Wang L, Aliferis CF. A comprehensive comparison of random forests and support vector machines for microarray-based cancer classification. BMC Bioinformatics. 2008;9:319. doi:10.1186/1471-2105-9-319

17. Tiwari AK. Machine Learning Based Approaches for Prediction of Parkinson’s Disease. Mach Learn Appl An Int J. 2016;3: 33–39. doi:10.5121/mlaij.2016.3203

18. Orhan U, Hekim M, Ozer M. EEG signals classification using the K-means clustering and a multilayer perceptron neural network model. Expert Syst Appl. Pergamon; 2011;38: 13475–13481. doi:10.1016/j.eswa.2011.04.149

19. Salem ABM, Revett K, Ei-Dahshan ESA. Machine learning in electrocardiogram diagnosis. Proc Int Multiconference Comput Sci Inf Technol IMCSIT’09. 2009;4: 429–433. doi:10.1109/IMCSIT.2009.5352689

20. Mena L, Gonzalez JA. Symbolic One-Class Learning From Imbalanced Datasets: Application in Medical Diagnosis. Int J Artif Intell Tools. 2009;18:273–309.doi:10.1142/S0218213009000135

21. Magoulas GD, Prentza A. Machine Learning in Medical Applications. Springer, Berlin, Heidelberg; 2001. pp. 300–307. doi:10.1007/3-540-44673-7_19

22. Itoh N, Hirashiki KI, Terada T, Kikuta M, Ishizuka SI, Koto T, et al. High sensitivity 900- MHz ISM band transceiver. IEICE Trans Fundam Electron Commun Comput Sci. 2005;E88–A: 498–505. doi:10.1109/72.159058

23. Yan H, Jiang Y, Zheng J, Peng C, Li Q. A multilayer perceptron-based medical decision support system for heart disease diagnosis. Expert Syst Appl. Pergamon; 2006;30:272–281. doi:10.1016/j.eswa.2005.07.022

24. Antal P, Fannes G, Timmerman D, Moreau Y, De Moor B. Bayesian applications of belief networks and multilayer perceptrons for ovarian tumor classification with rejection. Artif Intell Med. Elsevier; 2003;29:39–60. doi:10.1016/S0933-3657(03)00053-8

25. Morgan N, Bourlard H. Continuous speech recognition using multilayer perceptrons with hidden Markov models. International Conference on Acoustics, Speech, and Signal Processing. IEEE; pp. 413–416. doi:10.1109/ICASSP.1990.115720

26. Ren J, Shen X, Lin Z, Mech R, Foran DJ. Personalized Image Aesthetics. Proc IEEE Int Conf Comput Vis. 2017;2017–Octob: 638–647. doi:10.1109/ICCV.2017.76

27. Agarwal A, Negahban S, Wainwright MJ. Noisy matrix decomposition via convex relaxation: Optimal rates in high dimensions. Ann Stat. 2012;40:1171–1197. doi:10.1214/12-AOS1000

28. Zhang H, Gu C. Support Vector Machines versus Boosting. Dep Electr Eng Comput Sci Univ California, Berkeley, USA. 2006; 1--19. Available: http://citeseerx.ist.psu.edu/viewdoc/download?doi=10.1.1.117.343&rep=rep1&type=pdf

29. Bishop CM. Pattern Recognition and Machine Learning [Internet]. Jordan M, Kleinberg J, Scholkopf B, editors. Journal of Electronic Imaging. Springer; 2007. doi:10.1117/1.2819119

30. Kajdanowicz T, Kazienko P. Boosting-based multi-label classification. J Univers Comput Sci. Kluwer Academic Publishers; 2013;19: 502–520. doi:10.1023/A:1010933404324

31. Dreiseitl S, Ohno-Machado L. Logistic regression and artificial neural network classification models: A methodology review. J Biomed Inform. Academic Press; 2002;35:352–359. doi:10.1016/S1532-0464(03)00034-0

32. Martínková N, Nová P, Sablina O V., Graphodatsky AS, Zima J. Karyotypic relationships of the Tatra vole (Microtus tatricus). Folia Zool. 2004;53: 279–284. doi:10.1007/s13398-014-0173-7.2

33. Rubin DB. The Bayesian Bootstrap. Ann Stat. Institute of Mathematical Statistics; 1981;9: 130–134. doi:10.1214/aos/1176345338

34. Statnikov A, Aliferis CF, Tsamardinos I, Hardin D, Levy S. A comprehensive evaluation of multicategory classification methods for microarray gene expression cancer diagnosis. Bioinformatics. Oxford University Press; 2005;21:631–643. doi:10.1093/bioinformatics/bti033

35. Yeruva SLH, Paul Y, Oneal P, Nouraie M. Renal Failure in Sickle Cell Disease: Prevalence, Predictors of Disease, Mortality and Effect on Length of Hospital Stay. Hemoglobin. Taylor & Francis; 2016;40:295–299. doi:10.1080/03630269.2016.1224766

36. Al Khawaja SA, Ateya ZM, Al Hammam RA. Predictors of mortality in adults with Sickle cell disease admitted to intensive care unit in Bahrain. J Crit Care. W.B. Saunders; 2017;42: 238–242. doi:10.1016/j.jcrc.2017.07.032

